# Signatures of structural disorder in developing epithelial tissues

**DOI:** 10.1101/2024.02.12.579900

**Authors:** Christian Cupo, Cole Allan, Vikram Ailiani, Karen E. Kasza

## Abstract

Epithelial cells generate functional tissues in developing embryos through collective movements and shape changes. In some morphogenetic events, a tissue dramatically reorganizes its internal structure — often generating high degrees of structural disorder — to accomplish changes in tissue shape. However, the origins of structural disorder in epithelia and what roles it might play in morphogenesis are poorly understood. We study this question in the *Drosophila* germband epithelium, which undergoes dramatic changes in internal structure as cell rearrangements drive elongation of the embryo body axis. Using two order parameters that quantify volumetric and shear disorder, we show that structural disorder increases during body axis elongation and is strongly linked with specific developmental processes. Both disorder metrics begin to increase around the onset of axis elongation, but then plateau at values that are maintained throughout the process. Notably, the disorder plateau values for volumetric disorder are similar to those for random cell packings, suggesting this may reflect a limit on tissue behavior. In mutant embryos with disrupted external stresses from the ventral furrow, both disorder metrics reach wild-type maximum disorder values with a delay, correlating with delays in cell rearrangements. In contrast, in mutants with disrupted internal stresses and cell rearrangements, volumetric disorder is reduced compared to wild type, whereas shear disorder depends on specific external stress patterns. Together, these findings demonstrate that internal and external stresses both contribute to epithelial tissue disorder and suggest that the maximum values of disorder in a developing tissue reflect physical or biological limits on morphogenesis.

## I. INTRODUCTION

Epithelial tissue sheets comprise confluent packings of cells that undergo dramatic changes in overall shape and internal structure during development. The shapes and packings of cells in embryonic epithelia are diverse and can be regular and ordered or highly irregular and disordered. Moreover, the cell packings of a single tissue can change dramatically during development, mediated by changes in cell shapes and movements within the tissue [1–3]. For example, cells in the *Drosophila* germband epithelium initially tend to take on roughly regular, hexagonal shapes [4], which have shown to be optimized tilings for the densest arrangement of cells [5], but become more irregular as the tissue remodels during body axis elongation [4]. However, it is not well understood what biological factors regulate tissue disorder or how tissue disorder might influence tissue behavior and function.

Convergent extension of the *Drosophila* germband epithelium during body axis elongation is a powerful model for studying the biological and physical factors that regulate epithelial structure and remodeling [6–13]. Internal stresses from planar polarized actomyosin contractility and external stresses from neighboring developmental events both contribute to rapid elongation of the germband along the anterior-posterior (AP) axis of the embryo [7–12, 14–18]. This global tissue elongation is mediated by oriented cell rearrangements and cell shape changes that allow the tissue to remodel and flow [8, 10, 13–15, 19, 20]. Although it has been observed that cell shapes change and become more disordered during axis elongation [4, 13], the mechanisms driving this disorder and the resulting impacts on tissue elongation remain poorly understood.

The cell rearrangements and cell shapes changes that mediate tissue remodeling are thought to be driven by an interplay between local, internal stresses and global, embryo-scales stresses. Internal stresses are generated by a planar polarized pattern of actomyosin contractility, that is thought to arise from AP patterning systems [6, 7, 21] along with mechanosensitive feedback mechanisms [9, 18]. In addition, the germband experiences external stresses from ventral furrow formation (VFF) pulling along the dorsal-ventral (DV) axis and posterior midgut invagination (PMI) pulling along the AP axis [10– 12, 15, 16, 20, 22, 23]. A recent study has proposed that the stretch from the ventral furrow provides a mechanical cue to enrich myosin planar polarity and facilitate germband extension [18]. Together, these internal and external stresses are thought to control tissue structure and remodeling, but it has been difficult to decouple the effects of each.

Tissue structure is likely to influence biological and mechanical behaviors of the tissue. Work from our group and others has shown that the *average* cell shape in a tissue is a good predictor of whether the tissue will remodel and flow, similar to a fluid, or instead resist remodeling, similar to an elastic solid [13, 24–28]. Recent studies have demonstrated links between metrics for the *heterogeneity* in local tissue structure, i.e. structural disorder, and transitions in solid-like, liquid-like, and gas-like behavior of cultured tissues and developing embryos [29]. However, it remains unclear if and how structural order or disorder in a tissue might facilitate tissue remodeling.

Here, we study structural disorder in the developing *Drosophila* germband epithelium by analyzing volumetric and shear order parameters. We show that structural disorder in the germband increases during axis elongation in characteristic steps that coincide with developmental events in the embryo. We uncover that internal and external stress patterns contribute in distinct ways to volumetric and shear disorder, which are in turn linked with the rates and final extents of tissue remodeling. We find that the maximum magnitudes of structural disorder in the germband are in the range expected for purely random cell packings that have minimum cell-cell spacings, suggesting geometrical constraints on tissue disorder in this context. Overall, this work uncovers the biological and physical mechanisms that contribute to epithelial tissue disorder and highlight a central role for tissue disorder in epithelial remodeling and morphogenesis.

## II. RESULTS

### Structural disorder increases in the germband epithelium during body axis elongation

To explore how structural disorder evolves in developing epithelial tissues, we analyzed volumetric and shear disorder in the developing *Drosophila* embryo during body axis elongation. In *Drosophila*, the embryonic body axis rapidly doubles in length along the anterior-posterior (AP) axis through convergent extension movements of the germband epithelium (Fig. 1a) [6]. Tissue elongation is mediated in large part by rapid rearrangements of epithelial cells within the germband (Fig. 1b) [8, 19]. To analyze epithelial tissue structure, we fluorescently labeled cell membranes, imaged the embryo by confocal microscopy, segmented the timelapse movies, and triangulated the centroids of cells within the ventrolateral region of the germband (see Materials and Methods).

**FIG. 1.**
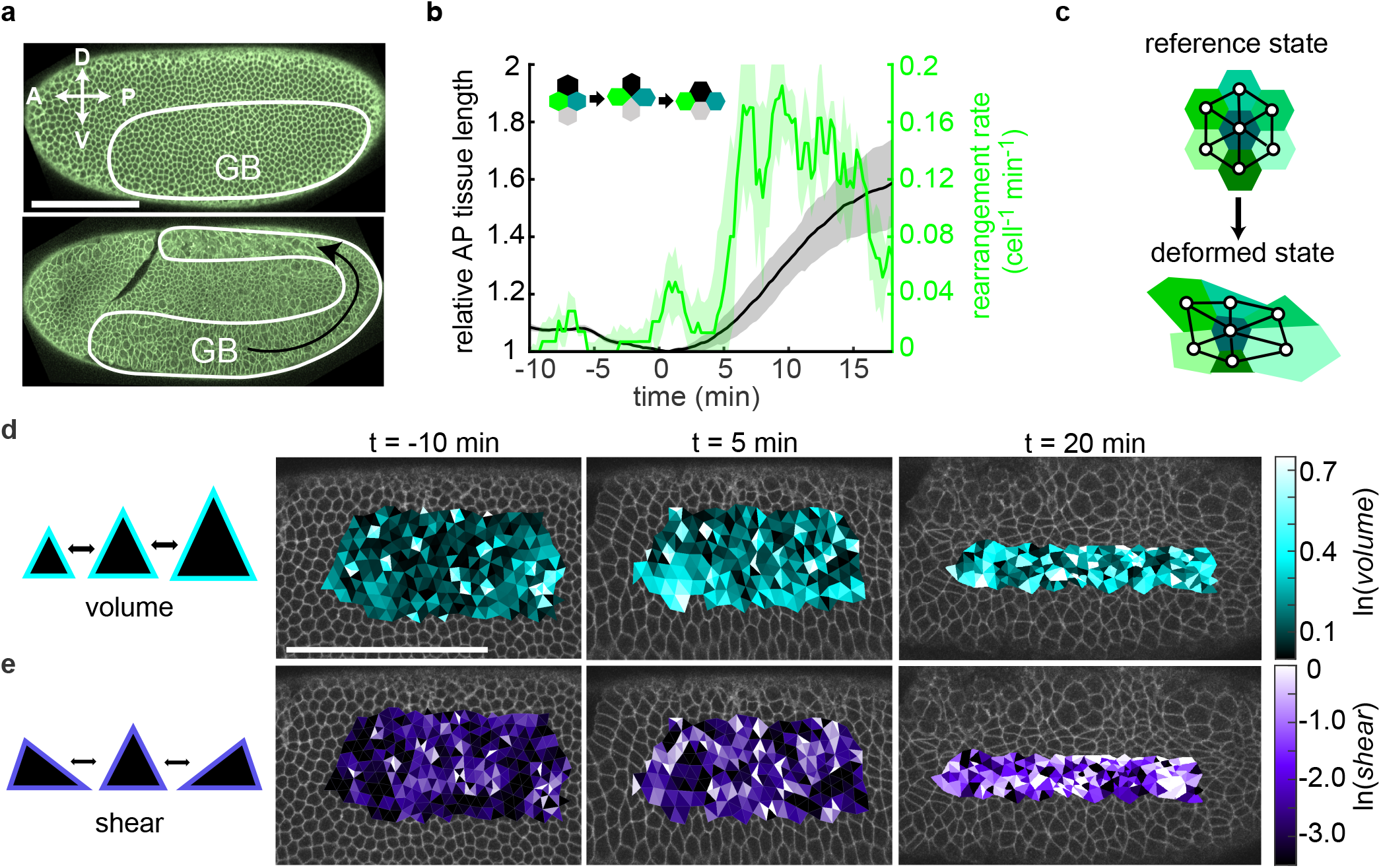
Structural disorder in the germband epithelium of the developing *Drosophila* embryo. (a) During body axis elongation, the germband epithelium (GB) undergoes convergent extension, narrowing along the dorsal-ventral (DV) axis and extending to twice its length along the anterior-posterior (AP) axis. (b) Tissue elongation (*black*) is facilitated by cellular rearrangements (*green*) that drive internal structural remodeling. The mean and standard deviation between 4 embryos is plotted. (c) Structural disorder in the tissue can be characterized by triangulating cell centroids and comparing to a reference configuration. (d,e) Triangulation of the germband at *t* = −10, 5, and 20 min. The color of each triangle represents its volumetric (density variation) or shear (irregularity compared to an equilateral triangle) deformation. Scale bar, 100 *µ*m.

To explore tissue structural disorder, we quantified the local volume and shear deformations in the triangulated tissue, as previously described [29]. Briefly, for the *n*th triangle in the tissue triangulation, the volume and shear invariants, *vol*_n_ and *shear*_n_, are calculated by comparing a deformed state to a reference state and determining the deformation (Fig. 1c). *vol*_n_ reflects how the triangle area deviates from the average area at each time point (Fig. 1d), and *shear*_n_ reflects the distortion from an equilateral triangle (Fig. 1e). Both invariants would be minimized if the triangulation comprises equilateral triangles of uniform area, i.e. a regular, isotropic honeycomb lattice has no volumetric or shear disorder. We find that the variation across the germband of *vol*_n_ (Fig. 1d) and *shear*_n_ (Fig. 1e) tends to increase over time during axis elongation.

To quantify the growing structural disorder, we calculated two structural order parameters, Φ_vol_ and Φ_shear_ (Materials and Methods) [29]. The volumetric order parameter, Φ_vol_ = ⟨[*ln*(*vol*_*n*_)]^2^⟩, represents density fluctu-ations across the tissue, and the shear order parameter, Φ_shear_ = ⟨ *ln*(*shear*_*n*_) ⟩, represents the irregularity of triangles. Prior to axis elongation, both order parameters take on small, relatively constant values (Fig. 2a-c), indicating low levels of structural disorder in the germband. Around the onset of axis elongation, Φ_vol_ and Φ_shear_ both begin to increase (Fig. 2a-c), indicating growing structural disorder associated with tissue elongation.

**FIG. 2.**
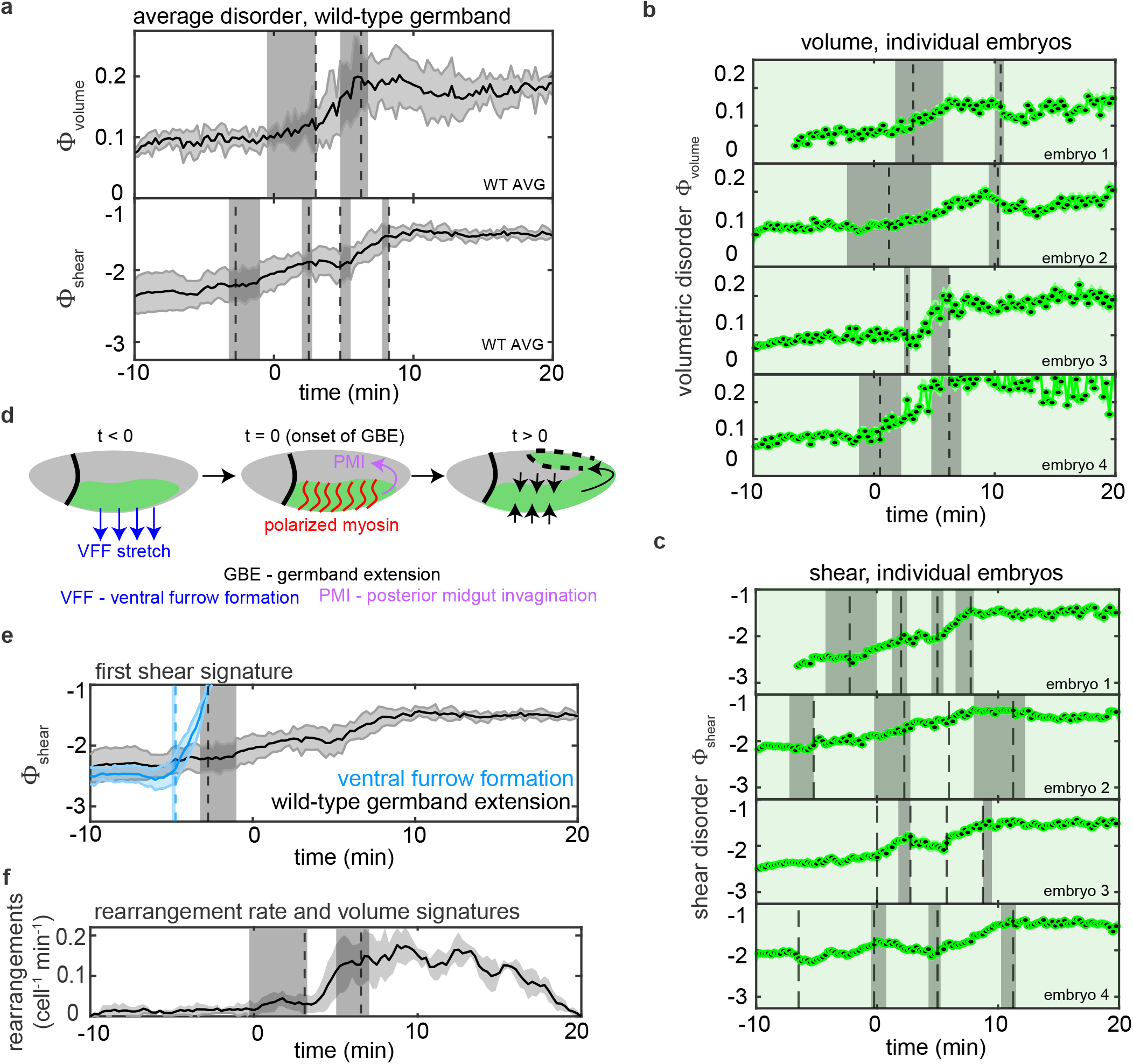
Structural disorder grows in the germband epithelium during axis elongation. (a) The volumetric order parameter Φ_vol_ and shear order parameter Φ_shear_ averaged across embryos (N=4 embryos). Shaded error bars represent standard deviation. Vertical dashed lines represent time points at which abrupt slope changes occur. The positions of the *n* changes in slope are identified using *n + 1* linear fits and minimizing the residuals. Vertical shaded regions around these lines represent values where the root mean squared deviation is within 10%. (b,c) Individual Φ_vol_ and Φ_shear_ curves over time for four wild-type embryos. Characteristic patterns or signatures of disorder are shared among embryos. (d) Schematics of axis elongation highlighting key developmental events in the germband (*green*) and neighboring regions of the embryo. (e) The first increase of Φ_shear_ in the germband (*black*) immediately follows an increase of Φ_shear_ in the neighboring ventral furrow region (*blue*). (f) The initial increase and final plateau in Φ_vol_ align with the onset of cell rearrangement events and the time point of maximum rearrangement rate, respectively.

We find that Φ_vol_ and Φ_shear_ increase in characteristic patterns that are consistent among wild-type embryos (Fig. 2a-c). We identified patterns in the growing disorder by finding points in the disorder vs. time curves where there are abrupt changes in slope. Overall, we find two volume and four shear disorder signatures. The first increase in disorder is an increase in Φ_shear_ 3 to 4 min before the onset of axis elongation, which occurs in the absence of changes in Φ_vol_ (Fig. 2a-c). Notably, this increase in germband shear disorder occurs just after the neighboring ventral furrow forms (Fig. 2d) and has its own increase in Φ_shear_ (Fig. 2e), suggesting that shear disorder in the germband is influenced by external stresses from the ventral furrow stretching the germband.

Just after the onset of axis elongation, which begins at *t* = 0 min, volumetric disorder begins to increase. The increase in Φ_vol_ aligns with the start of cell rearrangements (Fig. 2f), in which cells change contacts with their neighbors to promote tissue remodeling. These rearrangement events are driven by internal stresses from planar polarized actomyosin, suggesting that cell shape changes associated with rearrangement events contribute to volumetric disorder. Around the same time, Φ_shear_ temporarily plateaus, corresponding to tissue-scale reorientation of cell shapes from being stretched along the DV axis to being stretched along the AP axis, likely due to competing effects of the ventral furrow pulling along the DV axis and the posterior midgut pulling along the AP axis.

Notably, we find that both Φ_vol_ and Φ_shear_ plateau at *t* = 5 − 10 min at maximum values that are then maintained throughout the rest of the process. Φ_vol_ reaches a maximum 5 min after the onset of axis elongation, which corresponds to the timing of the maximum rate of cell rearrangement in the germband (Fig. 2f). Φ_shear_ plateaus a few minutes later. These disorder plateaus begin well before the end of axis elongation, while the tissue is still rapidly remodeling, suggesting that the maximum disorder plateaus might reflect a biological or physical constraint in the germband.

Taken together, these results demonstrate that structural disorder increases in the germband around the onset of axis elongation before reaching maximum plateau values that are then maintained through the rest of the process. Before reaching these plateaus, structural disorder increases in characteristic patterns or signatures, which are correlated with developmental events in the embryo.

### External stresses from the ventral furrow influ-ence germband structural disorder but not the maximum disorder reached

The timing of the initial increase in Φ_shear_ suggests a link between structural disorder in the germband and stresses from furrow formation in the neighboring ventral region of the embryo. To test this directly, we studied tissue disorder in *snail twist* (*sna twi*) mutant embryos in which ventral furrow formation (VFF) is disrupted (Fig. 3d). Initially, the germband of *sna twi* embryos displays lower values of Φ_vol_ and Φ_shear_ compared to wild-type embryos (Fig. 3a-c), indicating more ordered tissue structure. The initial increase in Φ_shear_ observed around *t* = −3 min in wild-type embryos is not present in *sna twi* embryos, demonstrating that this increase in shear disorder is driven by the ventral furrow. Instead, in *sna twi* embryos, Φ_shear_ and Φ_vol_ begin to increase just after *t* = 0 min.

**FIG. 3.**
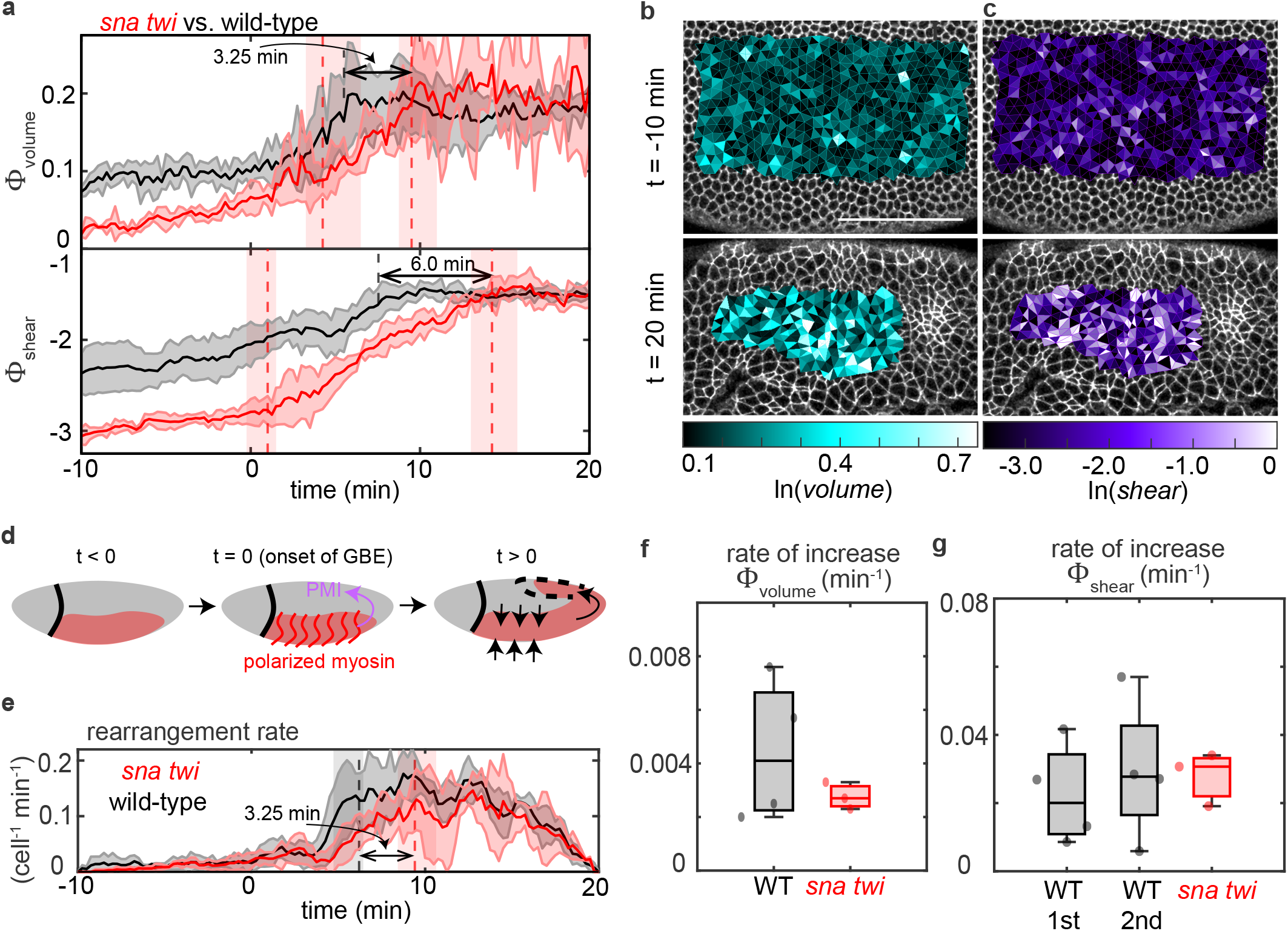
Structural disorder growth in the germband epithelium is delayed by disruption of ventral furrow invagination in *sna twi* mutant embryos. (a) Average Φ_vol_ and Φ_shear_ in the germband of *sna twi* mutant embryos (N = 3) (*red*) compared to wild type (N = 4) (*black*). Shaded error bars represent standard deviation between embryos. Vertical red dashed lines indicate positions of slope changes in curves from *sna twi* embryos. For reference, dashed, vertical black lines represent the timing of slope changes at the maximum disorder plateau in wild-type embryos. The maximum plateau values of Φ_vol_ and Φ_shear_ are delayed by 3.25 and 6 min, respectively, compared to wild-type embryos. (b,c) Volumetric (*cyan*) and shear (*purple*) deformation for the tissue triangulation at *t* = −10 and *t* = 20 min. (d) Schematics of key developmental events in *sna twi* mutant embryos with disrupted ventral furrow invagination. Disrupting the ventral furrow alters the external stress patterns acting on the germband. (e) Cell rearrangement rate in *sna twi* (*red*) and wild-type (*black*) embryos. Vertical dashed lines represent the slope change signature in Φ_vol_ that marks the time point of reaching the maximum Φ_vol_ in *sna twi* (*red*) and wild-type (*black*) embryos (c.f. panel a). In *sna twi* embryos, the delay in reaching maximum Φ_vol_ aligns with a delay in reaching the maximum rearrangement rate. (f) Rate of volumetric disorder growth (average slope of Φ_vol_(*t*) between slope changes) in wild-type and *sna twi* embryos. (g) Rate of shear disorder growth (average slope of Φ_shear_(*t*) between slope changes) in wild-type and *sna twi* embryos.

Despite starting out more ordered, Φ_shear_ and Φ_vol_ in the germband of *sna twi* embryos reach the same maximum plateau values as in wild-type embryos, but with a time delay (Fig. 3a). Once they begin to increase, the rates of disorder growth are comparable in *sna twi* and wild-type embryos (Fig. 3f,g), resulting in time delays of 3.25 min for Φ_vol_ and 6 min for Φ_shear_ for reaching the plateau values (Fig. 3a). *sna twi* embryos also display a delay in cell rearrangements, reaching the same maximum rearrangement rate as wild-type embryos but with a ∼3 min delay (Fig. 3e). These delays in *sna twi* embryos suggest that the ventral furrow might facilitate rapid axis elongation in wild-type embryos, an idea consistent with recent reports [18].

Notably, the maximum plateau values of Φ_shear_ and Φ_vol_ are the same in *sna twi* and wild-type embryos, indicating that external stresses from the ventral furrow do not affect the final structural disorder in the germband. One possible explanation is that loss of external stresses from the ventral furrow could be compensated for by other mechanisms. Alternatively, it is possible that the ventral furrow only influences germband tissue disorder during the first minutes of axis elongation before other factors dominate, such as internal myosin planar polarity (Fig. 3d). In either case, our results suggest that the maximum plateau values of structural disorder we observe in both wild-type and *sna twi* embryos reflect biological or physical constraints in the tissue that can be reached via different developmental trajectories.

### Internal stresses from myosin planar polarity play central roles in generating structural disorder in the germband

As external stresses from the ventral furrow were not sufficient to explain the structural disorder increases in the germband, we wondered if internal stresses associated with actomyosin contractility in the germband might be responsible for structural disorder in the tissue. To explore this possibility, we investigated two different mutants that disrupt AP patterning systems required for myosin planar polarity in the germband. *Krüppel* (*Kr*) mutants have partially disrupted AP patterning and display disrupted myosin planar polarity and reduced cell rearrangement in the central region of the germband (Fig. 4a). In contrast, *bicoid nanos torso-like* (*bcd nos tsl*) mutants have more fully disrupted AP patterning, displaying disrupted myosin planar polarity in the germband as well as disrupted posterior midgut invagination (Fig. 4b). Both mutants are expected to have disrupted internal stresses within the germband, while *bcd nos tsl* is expected to additionally have disrupted external stresses along the AP axis.

**FIG. 4.**
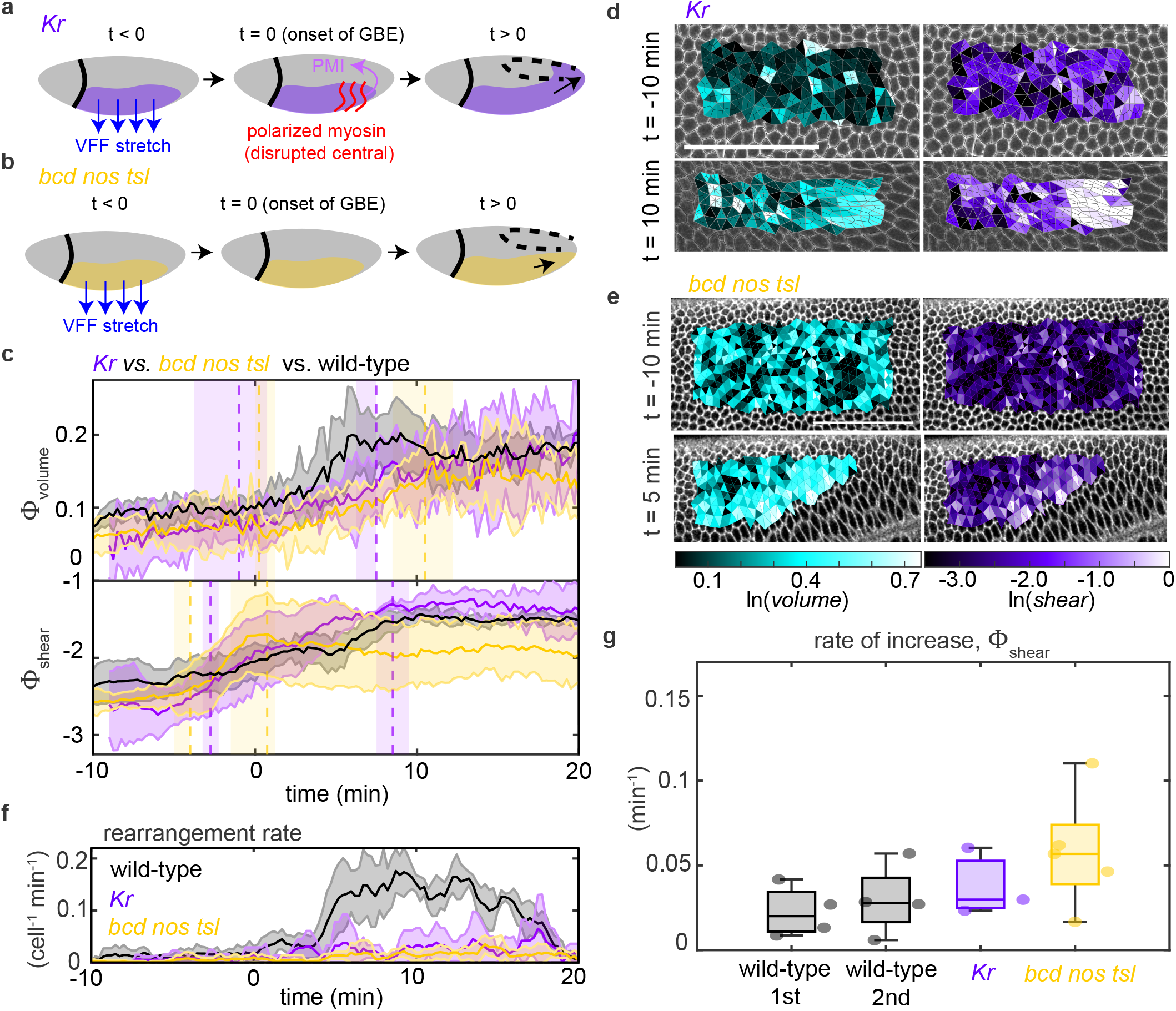
Structural disorder growth is disrupted in mutant embryos with disrupted AP patterning. (a) Schematics of key developmental events in *Kr* mutant embryos with disrupted myosin planar polarity in anterior and central regions of the germband. Disrupting myosin planar polarity alters the anisotropic internal stress patterns in the germband. (b) Schematics of key developmental events in *bcd nos tsl* mutant embryos with disrupted myosin planar polarity in the germband and disrupted posterior midgut invagination. Disrupting posterior midgut invagination alters the external stress patterns acting on the germband. (c) Average Φ_vol_ and Φ_shear_ in the germband of *Kr* (*purple*) (N = 3) and *bcd nos tsl* (*yellow*) (N = 5) mutants compared to wild type (*black*). Shaded error bars represent standard deviation between embryos. Vertical dashed lines indicate locations of slope changes within each embryo group. (d) Volumetric (*cyan*) and shear (*purple*) deformation for the tissue triangulation at *t* = −10 and 10 min in *Kr* mutants. (e) Volumetric (*cyan*) and shear (*purple*) deformation for the tissue triangulation in *bcd nos tsl* mutant embryos at *t* = −10 and 5 min. (f) Cell rearrangement rates in *Kr, bcd nos tsl*, and wild-type embryos. (g) Rate of shear disorder growth (average slope of Φ_shear_(*t*) between slope changes) for wild-type, *Kr*, and *bcd nos tsl* embryos.

Both mutants display low values of Φ_vol_ prior to axis elongation, similar to that in wild-type embryos (Fig. 4c). Φ_vol_ begins to increase around the same time as in wild-type embryos, just after *t* = 0, and increases at a slower rate, consistent with internal stresses and/or cell rearrangements driving this disorder. Notably, the *bcd nos tsl* mutant does not reach the maximum disorder values observed in wild-type embryos (Fig. 4c). The volumetric disorder reductions are associated with strongly reduced cell rearrangements in these embryos (Fig. 4f). These results show that volumetric disorder is strongly linked with AP patterning. The results from *Kr* mutants demonstrate that internal stresses and cell rearrangements associated with myosin planar polarity contribute to volumetric disorder. The stronger effects in *bcd nos tsl* mutants might be explained by more severely disrupted myosin planar polarity and/or disrupted posterior midgut invagination.

Both *Kr* and *bcd nos tsl* mutants also display low values of Φ_shear_ prior to axis elongation, similar to wild-type embryos, before beginning to increase around the onset of axis elongation (Fig. 4c). In *Kr* mutants, shear deformations are not evenly distributed across the germband, being more concentrated at the posterior end where midgut invagination is pulling on the germband (Fig. 4d). This effect is less pronounced in wild-type embryos in which cell rearrangements can help relax this stretch. *Kr* mutants reach about the same maximum plateau value of Φ_shear_ as wild-type embryos (Fig. 4c), suggesting that internal myosin planar polarity is not essential for generating high levels of shear disorder and that external stresses might play a more dominant role.

In *bcd nos tsl* mutants, Φ_shear_ increases rapidly just prior to the onset of axis elongation before plateauing early at a reduced level of shear disorder (Fig. 4c). Given the disruption of myosin planar polarity and posterior midgut invagination in these embryos, these results support the idea that this early increase in shear disorder is driven by the ventral furrow and then maintained throughout the rest of axis elongation. Interestingly, *bcd nos tsl* mutants appear to be more rapidly stretched by the ventral furrow compared to wild-type embryos, with a faster shear disorder increase (Fig. 4c,e,g), suggesting that the germband may be more compliant in *bcd nos tsl* mutants.

Taken together, these results demonstrate central roles for internal stresses associated with myosin planar polarity in volumetric disorder, although other factors also contribute, and highlight that external stresses associated with posterior midgut invagination and ventral furrow invagination contribute to shear disorder in the germband.

### Linking structural disorder to tissue elongation

To combine and visualize all of our data on structural disorder, we plotted the trajectories that wild-type and mutant embryos take in the Φ_vol_ and Φ_shear_ parameter space during axis elongation. We find that each embryo group takes a distinct path through the parameter space as volumetric and shear disorder grow. However, all trajectories appear to converge towards and not exceed the values of Φ_vol_≈ 0.20 and Φ_shear_≈ −1.5 (Fig. 5a), which correspond to the plateau values in wild type. It was unclear why volumetric and shear disorder plateau and what sets these plateau values.

**FIG. 5.**
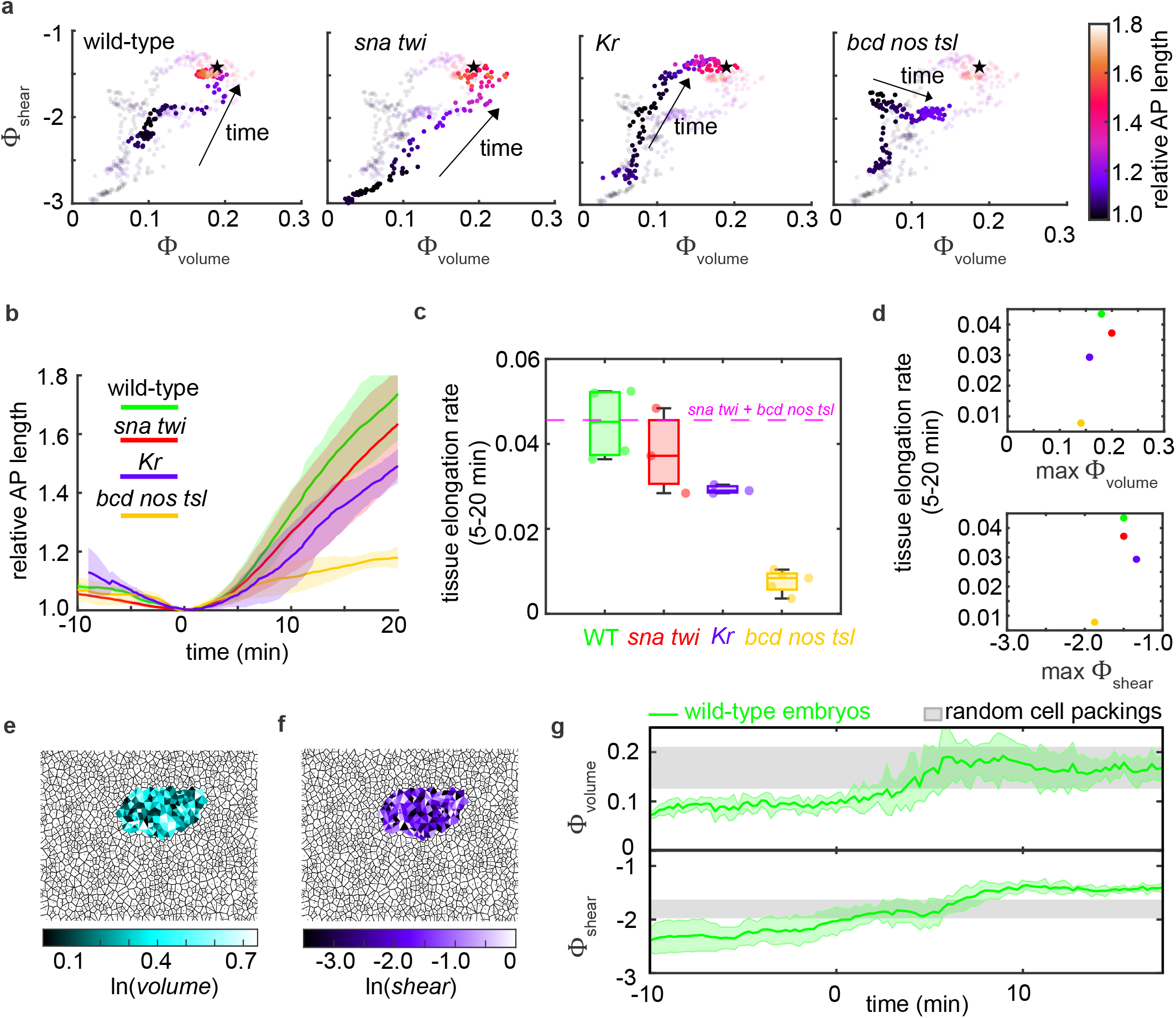
Linking volumetric and shear disorder patterns to tissue elongation. (a) Trajectories through the Φ_shear_ −Φ_vol_ parameter space over time during axis elongation. The extent of tissue elongation is indicated by the color of each data point. Stars, point towards which volumetric and shear disorder converge. (b) Tissue elongation over time for wild type (*green*), *sna twi* (*red*), *Kr* (*purple*), and *bcd nos tsl* (*yellow*). Mean *±*standard deviation between embryos. (c) Average tissue elongation rates between *t* = 5 min to *t* = 20 min for each group. The sum of elongation rates for *sna twi* and *bcd nos tsl* embryos is represented by the dashed magenta line. (d) Tissue elongation rate vs. plateau values of Φ_vol_ and Φ_shear_. (e,f) Randomly generated tilings of cell packings using densities and minimum spacings characteristic of the germband. Overlaid volumetric and shear deformations for the resulting triangulation. (g) The plateau value of Φ_vol_ in the wild-type germband (*green*) overlaps the range of values for random cell packings with densities and minimum spacings characteristic of the germband (*gray horizontal band*). The small, intermediate plateau of Φ_shear_ in the germband (*green*) overlaps with the range of values for random cell packings with densities and minimum spacings characteristic of the germband (*gray horizontal band*). The final plateau exceeds values for random packings, likely due to anisotropy in the germband.

One possibility is that tissue elongation requires a concomitant increase in disorder. In this case, structural disorder is essentially a readout of the history of tissue remodeling via cell rearrangements and/or cell stretching. However, this is not consistent with the observation that both Φ_vol_ and Φ_shear_ plateau at *t* = 5 −10 min in wild-type embryos, well before the tissue has fully elongated (Fig. 5b).

A second possibility is that structural disorder is instead a readout of the current rate of tissue remodeling. In this case, structural disorder should be directly related to the instantaneous rates of tissue elongation or cell rearrangement. The maximum value of disorder would then be set by embryonic mechanics that control tissue deformation rates. Supporting this idea, we found that Φ_vol_ reaches a maximum at the same time that the maximum rate of cell rearrangement is achieved in wild-type embryos at *t* = 5 min (c.f. Fig. 2a,f). However, after this time point, Φ_vol_ maintains its plateau value, while the cell rearrangement rate slowly decreases (c.f. Fig. 2a,f). Moreover, the average tissue elongation rates from *t* = 5 −20 min for wild type, *sna twi*, and *Kr* take on different values, while Φ_vol_ and Φ_shear_ have similar platea values (Fig. 5d). Thus, disorder is not simply related with either the extent or the instantaneous rate of tissue remodeling.

A third possibility is that the observed disorder plateaus might reflect some type of geometrical [30] or maximum entropy constraint [31]. In this case, disorder might increase due to tissue remodeling but then reach a maximum value set by limits on tissue disorder, even as the tissue continues remodeling. To explore this possibility, we needed a reference for comparing the magnitudes of Φ_vol_ and Φ_shear_ observed in the germband. We started by comparing our results to model tissues with random cell packings. We generated tilings of random cell packings using ranges of cell densities and minimum centroid spacings that we measured in the germband movies (Materials and Methods)(Fig. 5e,f) and quantified the range of values of Φ_vol_ and Φ_shear_ for the resulting packings (Fig. 5g).

Notably, the maximum plateau value of Φ_vol_ for the wild-type germband falls within the range we calculated for random packings with minimum cell centroid spacings characteristic of the germband (Fig. 5g). This indicates that, in terms of density fluctuations, cell packings in the germband are initially more ordered than random cell packings and become more disordered as the tissue remodels, up until having features similar to those of a random packing. This suggests that random packings with a minimum cell centroid spacing may represent a limit on tissue disorder in this context.

In contrast, Φ_shear_ in the germband reaches larger values than calculated for random cell packings (Fig. 5g). This is consistent with tissue-level cell stretching in the germband that can produce higher shear deformations than expected for random, isotropic packings. Interestingly, the values of Φ_shear_ for random packings do match the small, intermediate plateau we observe in wild-type embryos, during a period when cells in the tissue are reorienting (Fig. 5g). This suggests that random packings may also represent a limit on shear disorder in relatively isotropic tissues, and that tissue-level anisotropy from external or internal stresses can modulate this.

Finally, we asked how volumetric and shear disorder patterns are ultimately linked with the axis elongation process. Although tissue elongation rates were not simply related to either Φ_vol_ or Φ_shear_ (Fig. 5d), we noted that the embryos that elongated the most (wild type, *sna twi, Kr*) had *combinations* of relatively high Φ_vol_ and/or Φ_shear_, while the embryos that elongated significantly less (*bcd nos tsl*) had reduced values of both Φ_vol_ and Φ_shear_ (Fig. 5a-c). This suggests a combination of volumetric disorder, which we find to be linked with AP patterning and cell rearrangements, and shear disorder, which we find to be linked with external stress patterns, are central to tissue remodeling. Intriguingly, the sum of tissue elongation rates for *sna twi* (with disrupted ventral furrow) and *bcd nos tsl* (with disrupted myosin planar polarity and posterior midgut invagination) is nearly identical to the tissue elongation rate for wild type (Fig. 5c). This suggests that internal stresses associated with myosin planar polarity, external stresses from posterior midgut invagination, and external stresses from ventral furrow invagination may contribute in an additive fashion to axis elongation, although they are not additive in disorder. Taken together, these results demonstrate that rapid tissue elongation is linked with strong patterns of growing volumetric and shear disorder and suggest that there are physical limits on tissue disorder in this context.

## III. DISCUSSION

Here we showed that volumetric and shear structural disorder increase in the germband epithelium during body axis elongation in *Drosophila*. We find characteristic signatures of disorder that are shared among wild-type embryos and are correlated with developmental events involving changing patterns of internal and/or external stresses acting on the germband. By studying mutant embryos with altered patterns of internal and external stresses, we show that internal stress patterns associated with myosin planar polarity and cell rearrangement are strongly linked with volumetric disorder while external stress patterns associated with ventral furrow and posterior midgut invagination dominate shear disorder. Although the structural disorder metrics begin to increase around the onset of axis elongation, they both quickly plateau at values that are then maintained. These plateau values are similar to those for random cell packings, suggesting this may reflect a limit on tissue behavior. Finally, we find that high volumetric and shear disorder is linked with rapid tissue elongation, although the precise balance between volumetric and shear disorder is distinct in the germband of wild-type and mutant embryos that elongate significantly.

Our findings demonstrate that structural disorder grows during germband extension and is an important feature of this remodeling tissue. This extends prior work on topological disorder, which showed that the germband initially contains cells that are predominantly hexagonal in shape prior to axis elongation, before topological disorder increases during axis elongation [4]. In contrast, in the developing *Drosophila* wings and notum, cell rearrangements and junction fluctuations can increase geometric order, converting initially disordered cell packings into more hexagonally packed cells [32, 33]. In proliferating epithelia, cell divisions can also contribute to topological disorder during development [34]. Our results add to a growing body of work implicating geometric or structural signatures in jamming transitions, cell migration, wound healing, development, and cancer metastasis [13, 24–30, 35–39]. We find that the initial growth of structural disorder in the germband is correlated with the onset of rapid cell rearrangements and tissue remodeling, consistent with a possible solid-fluid-like transition [13]. However, the following nearly step-like disorder increases appear more like signatures of structural changes associated with specific developmental events.

It remains unclear if structural disorder is primarily an indicator of tissue remodeling or instead also feeds back to influence tissue remodeling. In particular, it is not known if the current structural order (disorder) in a tissue might facilitate (hinder) cell rearrangement and tissue remodeling. We find that after the germband reaches a maximum plateau value of volumetric disorder in wild-type and *sna twi* embryos, the cell rearrangement rates slowly begins to decrease. This suggests that cell rearrangement promotes disorder and that this disorder then might hinder further rearrangement. This is consistent with ideas from recent findings that disordered packings of cells extend more slowly than more ordered packings in some contexts [40, 41]. Future studies will be needed to fully address how tissue disorder impacts tissue remodeling and flow.

Our findings suggest that there are limits on the maximum structural disorder in the germband. The maximum volumetric disorder observed in the germband of wild-type and mutant embryos matches that of random tilings of cells with densities and minimum cell centroid spacings characteristic of the germband. In random cell packings, volumetric disorder is strongly influenced by the minimum spacing between cell centroids, suggesting that a minimum spacing between cells might underlie the limits on structural disorder we observe in the germband. This minimum spacing might arise because the apical domain is tightly regulated and maintained by epithelial cells or because nuclei might influence cell packings. Consistent with this possibility, recent reports highlight the importance of nucleus size and packings [42, 43].

Our findings support the notion that internal stresses from myosin planar polarity and external stresses from ventral furrow and posterior midgut invagination all contribute to tissue remodeling. There have been conflicting reports on the role of the ventral furrow in germband extension. It has recently been proposed that the stretch from ventral furrow invagination recruits myosin to cell junctions in the germband in a planar polarized fashion via mechanical feedback mechanisms, which then promote tissue flow [18]. However, other studies have shown that myosin planar polarity acts downstream of AP patterning systems [6, 7, 21] and have shown that the germband displays cell rearrangements and tissue elongation even when the ventral furrow is disrupted [10, 13, 44, 45]. Here, we find delays in disorder, cell rearrangement, and tissue elongation in *sna twi* mutant embryos, suggesting that the ventral furrow facilitates but is not required for tissue remodeling.

Taken together, our results identify growing structural disorder as a key feature of tissue remodeling. It remains unclear how structural disorder impacts tissue mechanics and might in turn influence patterns of cell rearrangements and tissue elongation during morphogenetic events. Moreover, it is not known how spatiotemporal patterns of cellular contractile and adhesive machineries drive structural disorder and might be influenced by tissue structure through mechanosensitive mechanisms. Future work will be needed to clarify these questions and explore how tissue disorder ultimately impacts tissue behavior and function.

## IV. MATERIALS AND METHODS

### Fly lines and genetics

Fly stocks were from the Bloomington Drosophila Stock Center (BDSC) unless otherwise noted. Wild-type control embryos were *yw* with two maternal copies of a sqh −gap43:mCherry transgene [46] to label cell membranes and two maternal copies of sqh −GFP:ZipWT to label myosin II [47] (not used in this study). The *Krüppel* (*Kr*) mutant fly stock was a gift of Eric Wieschaus [6]. *Kr* embryos were zygotic *Kr* ^1^ mutants with a sqh −gap43:mCherry transgene. The movies of *snail twist* (*sna twi*) and *bicoid nanos torso-like* (*bcd nos tsl*) mutant embryos were from Wang., et al. [13]. *sna twi* embryos were zygotic *sna*^*IIG*05^ *twi* ^*DfS*60^ mutants and expressed Spider:GFP to visualize cell outlines. The *bcd nos tsl* maternal mutant embryos were the progeny of *bcd* ^*E*1^ *nos*^*L*7^ *tsl* ^146^ homozygous females and expressed Resille:GFP to visualize cell outlines.

### Embryo preparation

Embryos were generated at 23?. Embryos were collected at stage 6, dechorionated in 50% bleach for 2 min, washed with water for 2 min, and mounted on a custom-made imaging chamber in a 50:50 mixture of halocarbon oils 27 and 700 between an oxygen-permeable membrane (YSI Incorporated) and a glass coverslip.

### Confocal time-lapse imaging

Imaging was performed at room temperature on a Zeiss LSM 880 laser-scanning confocal microscope. Embryos were placed in a ventrolateral or ventral orientation for imaging. A 561 nm diode laser was used for imaging mCherry, and a 488 nm Argon laser excitation was used for GFP. Imaging was performed using the Airyscan FAST module and a C-Apo 40X/1.2 NA water-immersion objective (Carl Zeiss, Germany). Movies were acquired as 9.1 *µ*m z-stacks with a 0.7 *µ*m z-step every 15 seconds. The first image was captured at the most apical surface of the embryo. Movies were rotated and z-projections taken using the Fiji distribution of ImageJ [48, 49]. Each z-projection was performed by taking a maximum intensity projection of 3 consecutive z-slices at the apical junctional plane of the tissue.

Separate movies were captured for imaging the germband and ventral furrow regions of the embryo. Embryos for germband movies were oriented ventrolaterally, and embryos for ventral furrow movies were oriented ventrally. Regions of interests for each movie were defined, and cells were segmented based on cell membrane markers. For the ventral furrow movies, analysis concludes prior to tissue invagination and focuses on in-plane stretching.

### Cell segmentation analysis

Cell outlines were identified using a custom MATLAB code that implements a watershed algorithm [50] combined with a mask. The segmentation output was imported into the SEGGA tissue segmentation and analysis software [44] using a custom MATLAB script, and manual corrections were performed using the SEGGA GUI. In the final segmentation, each cell is represented as a polygon with straight edges. Custom MATLAB scripts were used to further process and analyze data.

### Aligning movies

To compare movies, *t = 0* min was taken as the onset of germband extension, which was identified by finding the time point corresponding to the minimum germband tissue length, when considering the group of cells that were in the ROI at all time frames.

### Rearrangement rate analysis

Cells within the ROI were tracked and cell rearrangement events identified using SEGGA [44]. The number of rearrangements per cell in each time frame was calculated by dividing the number of cell neighbors lost in each time frame by the number of cells analyzed in that frame.

### Calculation of Φ_vol_ and Φ_shear_ and identifying transitions in these disorder metrics

The tissue was triangulated by finding the centers of all cells in the ROI and connecting adjacent centers. If four or more segments from a grouping of cells meet at a single node, an additional node was added at this point to create the triangulation. Using this triangulation, Φ_vol_ and Φ_shear_ were calculated as shown previously [29]. Transitions in Φ_vol_(*t*) and Φ_shear_(*t*) were identified by first visually identifying the number of slope changes (*n*) and then using a custom MATLAB script to fit (*n + 1*) linear fits to the data until the residuals were minimized. Errorbars for the timing of the transitions correspond to where the root mean squared deviation was within 10%.

### Random cell tiling simulation

We generated randomly organized cell packings and calculated Φ_vol_ and Φ_shear_ for these random packings. To do this, we seeded points within a known area to act as the nucleation centers, i.e. random sequential addition (RSA). The number of nucleation points was varied to match the range of cell densities observed in germband movies. Additionally, we varied the minimum distance between nucleation points to match the minimum spacing of cell centers observed in germband movies of 3-4 microns; this may help to account for biological mechanisms that maintain cell size and shape and/or incompressibility of cell nuclei or other internal cellular structures. We generated five unique Voronoi tilings and moved the seed points to the center of mass for each Voronoi cell (centroidal Voronoi diagram algorithm) [51]. We did this for each set of parameters and analyzed 5 different ROIs for each image. Each image was processed with the same pipeline as the germband movies. Briefly, the images were segmented with our algorithm and manually corrected in SEGGA before calculating Φ_vol_ and Φ_shear_.

### Error bars

Data plotted represent mean value ± standard deviation unless otherwise noted.

## Data availability

The data in this paper is available from the corresponding author upon reasonable request.

## Code availability

Custom MATLAB scripts are available from the corresponding author upon reasonable request.

## Acknowledgements

We thank Erika Kusaka for help generating fly stocks, Rob Campbell for the MATLAB function shadederrorbar, and members of the Kasza Lab, Andrew Cupo, Haiqian Yang, Ming Guo, and Max Bi for helpful discussions. This work was supported by the National Institute of General Medical Sciences of the National Institutes of Health Award Number R35GM138380 to K.E.K. and Eunice Kennedy Shriver National Institute of Child Health & Development of the National Institutes of Health Grant F31HD105405 to C.C. The content is solely the responsibility of the authors and does not necessarily represent the official views of the National Institute of Health. K.E.K. holds an NSF CAREER Award, Packard Fellowship, and Sloan Research Fellowship in Physics.

## Author Contributions

C.C. and K.E.K. initiated the project, designed the experiments, and developed the project. C.C performed all experiments. C.C. and C.A. developed the code for image processing and data analysis. C.C. and V.A. created the plots, images, and graphics for the figures. C.C. and K.E.K. wrote the manuscript. All authors contributed to data interpretation and approved the final manuscript.

